# Drug repurposing screening identified tropifexor as a SARS-CoV-2 papain-like protease inhibitor

**DOI:** 10.1101/2021.12.02.471030

**Authors:** Chunlong Ma, Yuyin Wang, Juliana Choza, Jun Wang

**Author notes:** **Corresponding Author Jun Wang** − Department of Pharmacology and Toxicology, College of Pharmacy, The University of Arizona, Tucson, Arizona 85721, United States.

## Abstract

The global COVID-19 pandemic underscores the dire need of effective antivirals. Encouraging progress has been made in developing small molecule inhibitors targeting the SARS-CoV-2 RNA-dependent RNA polymerase (RdRp) and main protease (M^pro^). However, the development of papain-like protease (PL^pro^) inhibitors faces several obstacles. Nevertheless, PL^pro^ represents a high-profile drug target given its multifaceted roles in viral replication. PL^pro^ is involved in not only the cleavage of viral polyprotein but also modulation of host immune response. In this study, we conducted a drug-repurposing screening of PL^pro^ against the MedChemExpress bioactive compound library and identified three hits, EACC, KY-226, and tropifexor, as potent PL^pro^ inhibitors with IC_50_ values ranging from 3.39 to 8.28 µM. The three hits showed dose-dependent binding to PL^pro^ in the thermal shift assay. In addition, tropifexor inhibited the cellular PL^pro^ activity in the FlipGFP assay with an IC_50_ of 10.6 µM. Gratifyingly, tropifexor showed antiviral activity against SARS-CoV-2 in Calu-3 cells with an EC_50_ of 4.03 µM, a 7.8-fold increase compared to GRL0617 (EC_50_ = 31.4 µM). Overall, tropifexor represents a novel PL^pro^ inhibitor that can be further developed as SARS-CoV-2 antivirals.

---

The etiological agent of COVID-19 is SARS-CoV-2, a single-stranded, positive-sense RNA virus that belong to the β-coronavirus genera. Given the catastrophic impact of COVID-19 on public health and global economy, researchers around of the globe are working relentlessly to develop vaccines and antiviral drugs. This effort led to the approval of vaccines and antiviral drugs in record breaking speed. Two mRNA vaccines from Moderna and Pfizer, and one adenovirus vaccine from Johnson and Johnson were approved by FDA.^1^

Although vaccines are the mainstay in combating the pandemic, antiviral drugs are nevertheless needed as complementary strategies. Vaccines are preventative, while antiviral drugs can be used for the treatment of COVID patients. In addition, the mRNA vaccines target the viral spike protein, which is prone to mutation as shown by the variants of concerns including the Delta variant and the most recent Omicron variant.^2^ As a result, vaccines might need to be frequently updated to match the circulating strains. In comparison, small molecule antiviral drugs targeting the conserved viral proteins are expected to have broad-spectrum antiviral activity and a high genetic barrier to drug resistance. The viral RNA-dependent RNA polymerase (RdRp) inhibitor remdesivir is the first FDA-approved COVID drug.^3^ In addition, the second RdRp inhibitor molnupirivir^4-6^ and the main protease (M^pro^) inhibitor PF-07321332 (Paxlovid)^7^ are likely to become the first oral COVID drugs.

Despite the encouraging progress, additional antiviral drugs with a novel mechanism of action are still in dire need to override the emergence of new mutations. They can be used either alone or in combination with existing RdRp inhibitors or M^pro^ inhibitors to combat not only current COVID-19 pandemic, but also future coronavirus outbreaks.

SARS-CoV-2 expresses two viral proteases, the M^pro^ and papain-like protease (PL^pro^), during viral replication. Both M^pro^ and PL^pro^ are cysteine proteases that mediate the cleavage of viral polyprotein during viral replication.^8^ In addition, PL^pro^ desregulates the host immune responses by cleaving ubiquitin and interferon-stimulated gene 15 protein (ISG15) from host proteins.^9^ Therefore, inhibiting PL^pro^ is a two-pronged approach in protecting host cells from viral infection.

PL^pro^ is a 35-KDa domain of Nsp3, a 215-KDa multidomain protein that is a key component of the viral replication complex.^10^ Compared to PL^pro^ from SARS-CoV, SARS-CoV-2 PL^pro^ displays decreased deubiquitination activity and enhanced deISGlyation activity.^9, 11^In contrast to M^pro^, PL^pro^ is a more challenging drug target mainly for two reasons. First, the protein substrate of PL^pro^ consists of LXGG.^12^ Accordingly, there is a lack of drug binding pockets in the S1 and S2 subsites. As such, majority of reported PL^pro^ inhibitors are non-covalent inhibitors that bind to the S3 and S4 subsites that are located more than 10 Å away from the catalytic cysteine C111.^13-15^ Second, PL^pro^ bears structural similarities to human deubiquitinases and delSGylases,^16^ which presents a challenge in developing selective PL^pro^ inhibitors. Despite extensive high-throughput screening and lead optimization,^11, 13-15, 17-18^ GRL0617 and its analogs remain the most potent PL^pro^ inhibitors reported so far. To identify structurally novel PL^pro^ inhibitors, we conducted a drug repurposing screening and identified EACC, KY-226, and tropifexor, as potent PL^pro^ inhibitors with IC_50_ values ranging from 3.39 to 8.28 µM. Their mechanism of action was further characterized in the thermal shift binding assay and the FlipGFP protease assay. Gratifyingly, tropifexor also had potent antiviral activity against SARS-CoV-2 in Calu-3 cells with an EC_50_ of 4.03 µM. Overall, tropifexor represents a potent PL^pro^ inhibitor with a novel scaffold that can be further developed as SARS-CoV-2 antivirals.

## RESULTS AND DISCUSSION

### High-throughput screening of SARS-CoV-2 PL^pro^ inhibitors

Using the previously optimized FRET assay condition,^15^ we performed a high-throughput screening of SARS-CoV-2 PL^pro^ against the MedChemExpress bioactive compound library which consists of 9,791 compounds including FDA-approved drugs, clinical candidates, and natural products. All compounds were originally screened at 40 µM, and hits showing more than 50% inhibition were further titrated to determine the IC_50_ values. GRL0617 was included as a positive control. In total, three compounds, EACC, KY-226, and tropifexor (Figure 1A), were identified as positive hits with IC_50_ values of 8.28, 3.39, and 5.11 µM, respectively (Figure 1B). In comparison, the IC_50_ value for the positive control GRL0617 was 1.66 µM (Figure 1B). Next, the broad-spectrum activity of the three hits was tested against SARS-CoV PL^pro^ (Figure 1C) and MERS-CoV PL^pro^ (Figure 1D). It was found that EACC, KY-226, and tropifexor retained potent inhibition against SARS-CoV PL^pro^ with IC_50_ values of 6.28, 3.53, and 5.54 µM, respectively (Figure 1C). In contrast, EACC and KY-226 were weak inhibitors of MERS-CoV PL^pro^ with IC_50_ values of 27.8 and 30.6 µM, while GRL0617 was inactive (IC_50_ > 60 µM) (Figure 1D). Nevertheless, tropifexor showed higher potency against MERS-CoV PL^pro^ with an IC_50_ of 2.32 µM (Figure 1D). The hits were further counter screened against the SARS-CoV-2 M^pro^ to rule out promiscuous cysteine protease inhibitors.^19-22^ It was found that EACC and KY-226 were not active (IC_50_ ≥ 60 µM), while tropifexor had weak inhibition with an IC_50_ of 43.65 µM, which corresponds to a selectivity index (SI) of 8.5 (Figure 1E). These results suggest the inhibition of SARS-CoV-2 PL^pro^ by tropifexor is specific. The inhibition of PL^pro^’s deubiquitination and deISGlyation activities were characterized using the Ub-AMC and ISG15-AMC substrates, respectively.^14-15^ While EACC and KY-226 were inactive in inhibiting the deubiquitinase activity of PL^pro^ (IC_50_ > 100 µM), tropifexor showed moderate activity with an IC_50_ of 18.85 µM (Figure 1F). Similarly, EACC and KY-226 were not active in inhibiting the deISGlyation activity of PL^pro^ (IC_50_ > 80 µM), tropifexor showed does-dependent inhibition with an IC_50_ of 27.22 µM (Figure 1G). Overall, tropifexor appears to be the most promising hit with consistent inhibition against SARS-CoV-2, SARS-CoV, and MERS-CoV PL^pro^s. In addition, tropifexor also inhibited the deubiquitination and deISGlyation activities of SARS-CoV-2 PL^pro^, albeit at lower potency.

**Figure 1.**
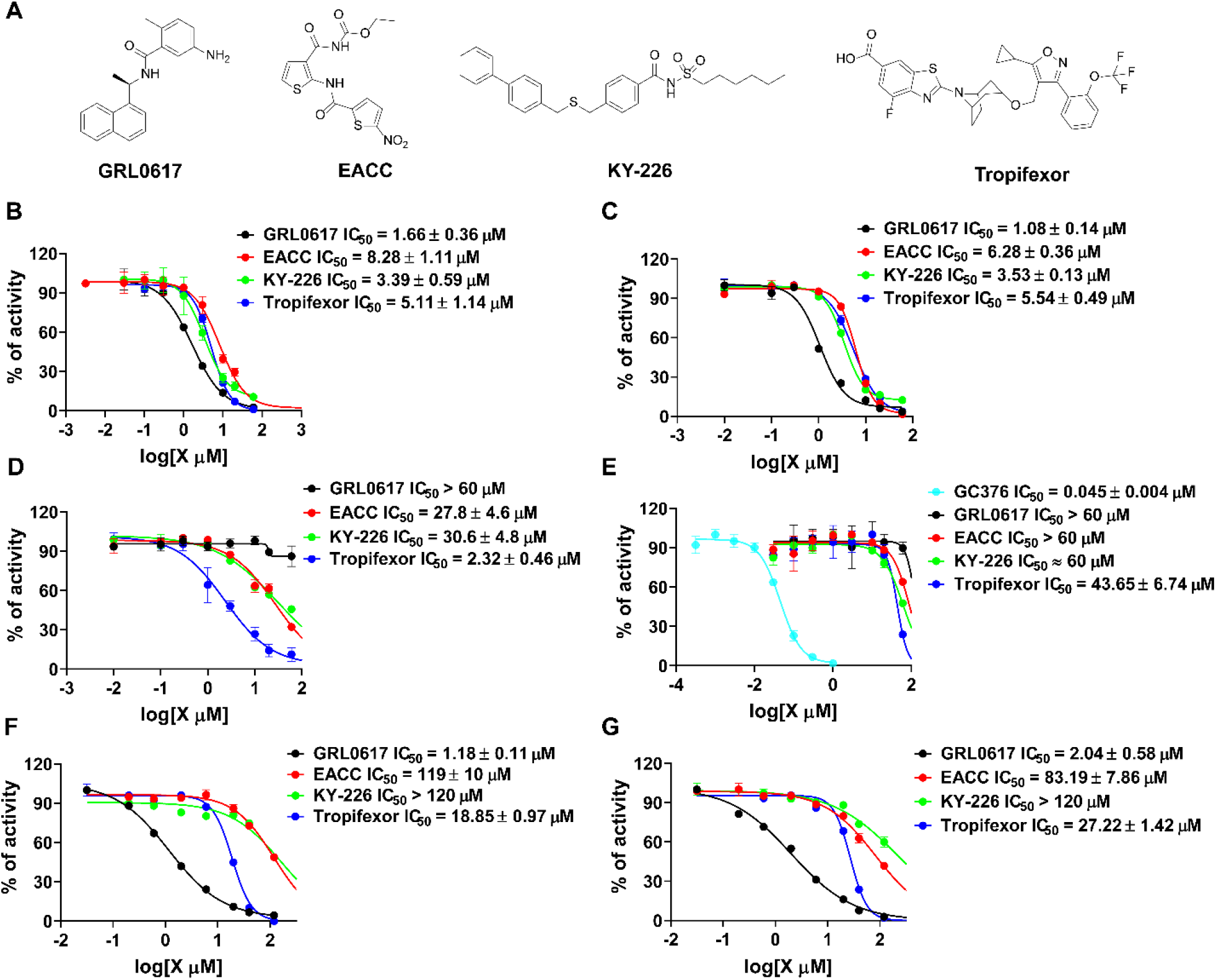
Characterization of SARS-CoV-2 PL^pro^ inhibitors identified from the high-throughput screening. (A) Chemical structures of the positive control GRL0617 and the three hits EACC, KY-226, and tropifexor. (B) IC_50_ curves of the hits in inhibiting SARS CoV-2 PL^pro^ with the FRET peptide substrate 1. (C) IC_50_ curves of the hits in inhibiting SARS CoV PL^pro^ with the FRET peptide substrate 1. (D) IC_50_ curves of the hits in inhibiting MERS-CoV PL^pro^ with the FRET peptide substrate 1. (E) IC_50_ curves of the hits in inhibiting SARS CoV-2 M^pro^ with the FRET peptide substrate 2. (F) IC_50_ curves of the hits in inhibiting SARS CoV-2 PL^pro^ with the Ub-AMC substrate. (G) IC_50_ curves of the hits in inhibiting SARS CoV-2 PL^pro^ with the ISG15-AMC substrate. Please refer to the methods and materials section for assay conditions. Values represent the average ± standard deviation of three replicates.

### Pharmacological characterization of the hits in the thermal shift binding assay and the cell-based FlipGFP PL^pro^ assay

The mechanism of action of EACC, KY-226, and tropifexor in inhibiting SARS-CoV-2 PL^pro^ was further characterized in the thermal shift assay and the cell-based FlipGFP PL^pro^ assay.^15, 19^-^20, 23^ Similar to the positive control GRL0617, all three hits displayed dose-dependent binding to PL^pro^ as revealed by the enhanced melting temperatures with increasing drug concentration (Figure 2).

**Figure 2.**
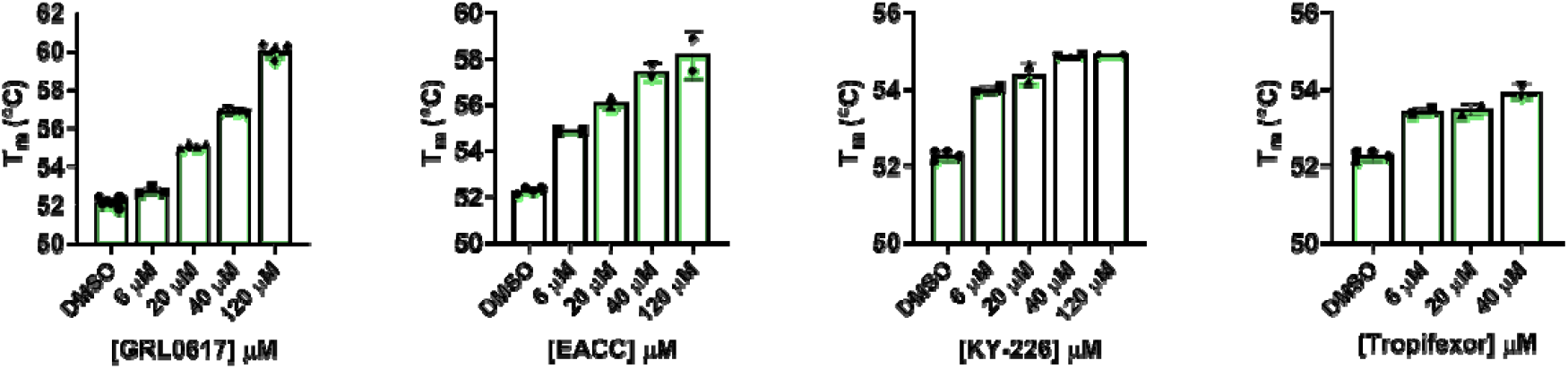
Thermal shift binding assay of SARS-CoV-2 PL^pro^ protease against identified inhibitors. All inhibitors display dose-dependent melting temperature (T_m_) shift. Values represent the average ± standard deviation of three replicates.

Next, we tested the three hits in the FlipGFP PL^pro^ assay.^15, 19-20^ The FlipGFP PL^pro^ was recently developed by us as a surrogate assay to quantify the cellular activity of PL^pro^ inhibitors in the biological safety level 2 facility, and we have shown that there is positive correlation between the FlipGFP IC_50_ values with the SARS-CoV-2 antiviral EC_50_ values.^15^ The FlipGFP assay is a virus free cell-based protease assay in which the 293T cells were transfected with PL^pro^ and the GFP reporter. The GFP reporter consists of two fragments,^24-25^ the β1-9 template, and the β10-11 strands that are constrained in the parallel inactive conformation through a PL^pro^ substrate linker. Upon cleavage of the substrate linker, the β10 and β11 strands become parallel and can associate with the β1-9 template, leading to increased GFP signal. mCherry is included as an internal control to normalize transfection efficacy and compound cytotoxicity. In principle, the normalized GFP/mCherry ratio is proportional to the enzymatic activity of PL^pro^. The advantages of FlipGFP assay compared to the FRET assay is that it can rule out compounds that are cytotoxic, membrane impermeable, and having off-target effects that prevent cellular on-target engagement.^19-20^

In the FlipGFP assay, the positive control GRL0617 showed dose-dependent inhibition with an IC_50_ of 14.67 µM, while the negative control GC376 was not active (IC_50_ > 60 µM) (Figure 3A, B). The results from EACC and KY-226 were not conclusive due to the cytotoxicity of the compounds. Tropifexor had an IC_50_ of 10.60 µM, but a low selectivity index (CC_50_ = 29.77 µM, SI = 2.8) (Figure 3A, B). In summary, the FlipGFP assay results suggest tropifexor might have antiviral activity against SARS-CoV-2.

**Figure 3.**
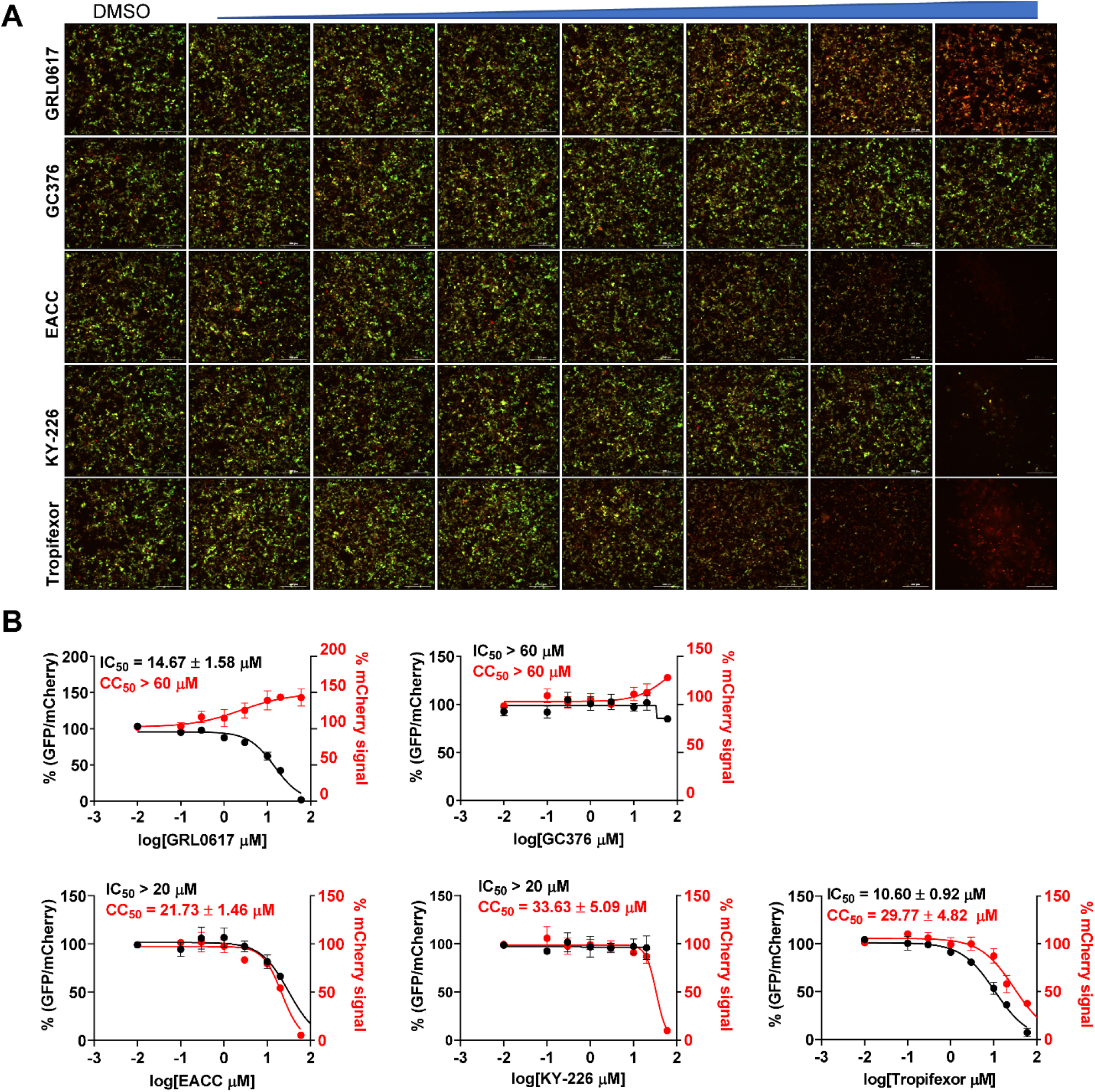
FlipGFP SARS CoV-2 PL^pro^ assay to determine cellular protease inhibitory activity of identified inhibitors. (A) Representative images of FlipGFP-PL^pro^ assay with increasing concentrations of GRL0617 (positive control), GC376 (negative control), EACC, KY-226, and tropifexor. GRL0617 showed does-dependent decrease of GFP signal with the increasing drug concentration, while almost no GFP signal change was observed with the increasing concentration of negative control compound GC376. (B) Dose-response curves of the GFP/mCherry ratio with increasing drug concentrations. mCherry signal alone was used to calculate transfection efficiency and compound cytotoxicity. All three hits displayed significant cytotoxicity at high drug concentrations. Values represent the average ± standard deviation of three replicates.

### Antiviral activity of hits against SARS-CoV-2 in Calu-3 cells

The antiviral activity of EACC, KY-226, and tropifexor in inhibiting SARS-CoV-2 replication in Calu-3 cells was tested using the immunofluorescence assay (Figure 4). Calu-3 is TMPRSS2-positive and is a close mimetic of the human respiratory epithelial cells,^26^ enabling it a widely accepted cell line for SARS-CoV-2 studies.^19, 27^ The positive control GRL0617 had an EC_50_ of 31.4 µM (Figure 4A). EACC did not show antiviral activity at non-toxic drug concentration (EC_50_ > 35 µM, CC_50_ = 35.29 µM) (Figure 4B).

**Figure 4.**
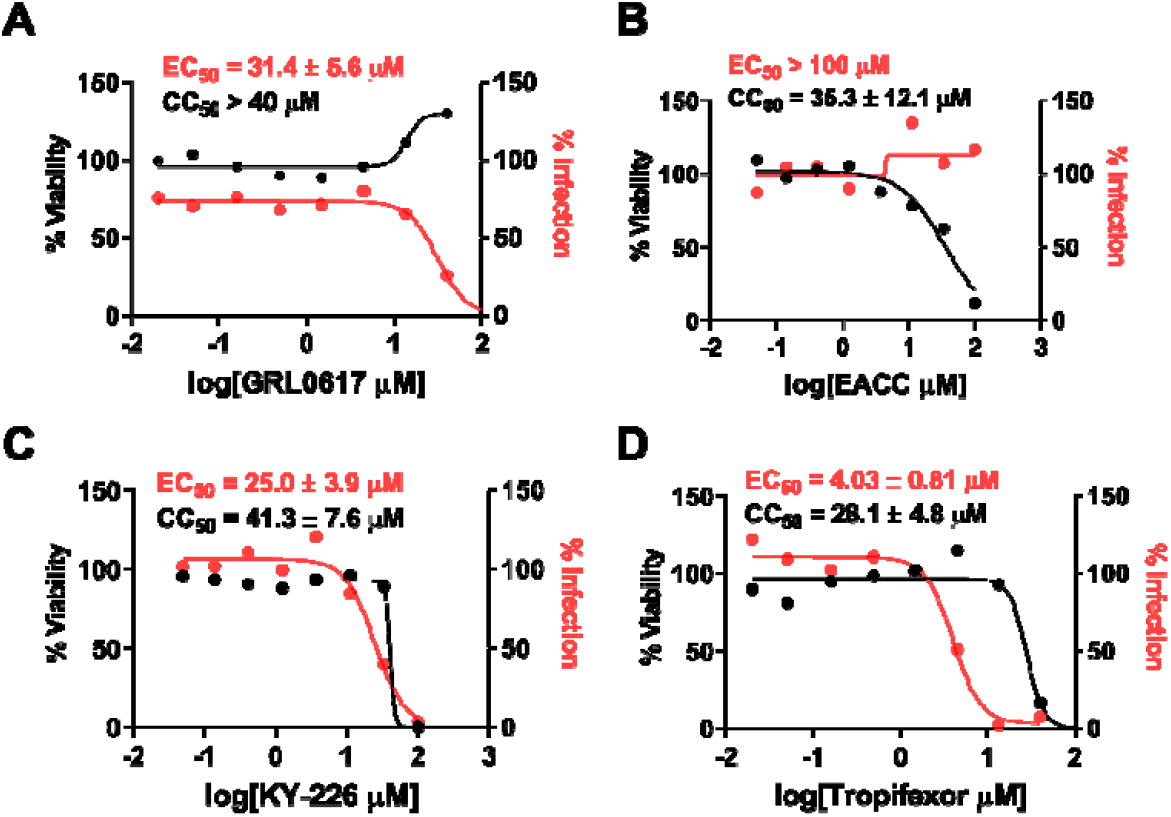
Antiviral activity of SARS-CoV-2 PL^pro^ inhibitors GRL0617 (A), EACC(B), KY-226 (C), and tropifexor (D) against SARS-CoV-2 in Calu-3 cells. The results were quantified by immunofluorescence assay. Values represent the average ± standard deviation of three replicates.

Gratifyingly, both KY-226 and tropifexor had improved antiviral activity against SARS-CoV-2 with EC_50_ values of 25.0 (Figure 4C) and 4.03 µM (Figure 4D), respectively.

While KY-226 had a low selectivity index (SI = 1.65), tropifexor had a moderate selectivity window (SI = 6.97) and the observed antiviral activity was clearly not caused by the cytotoxicity of the compound.

### Molecular docking of EACC, KY-226, and tropifexor in SARS-CoV-2 PL^pro^

To gain insights of the binding mode of the three hits, we performed molecular docking with Schrödinger Glide using the wild-type SARS-CoV-2 PL^pro^ structure we recently solved (PDB: 7JRN).^15^ EACC, KY-226 and tropifexor all fit snuggly into the U-shape binding pocket that is covered by the BL2 loop where GRL0617 binds (Figure 5A).

**Figure 5.**
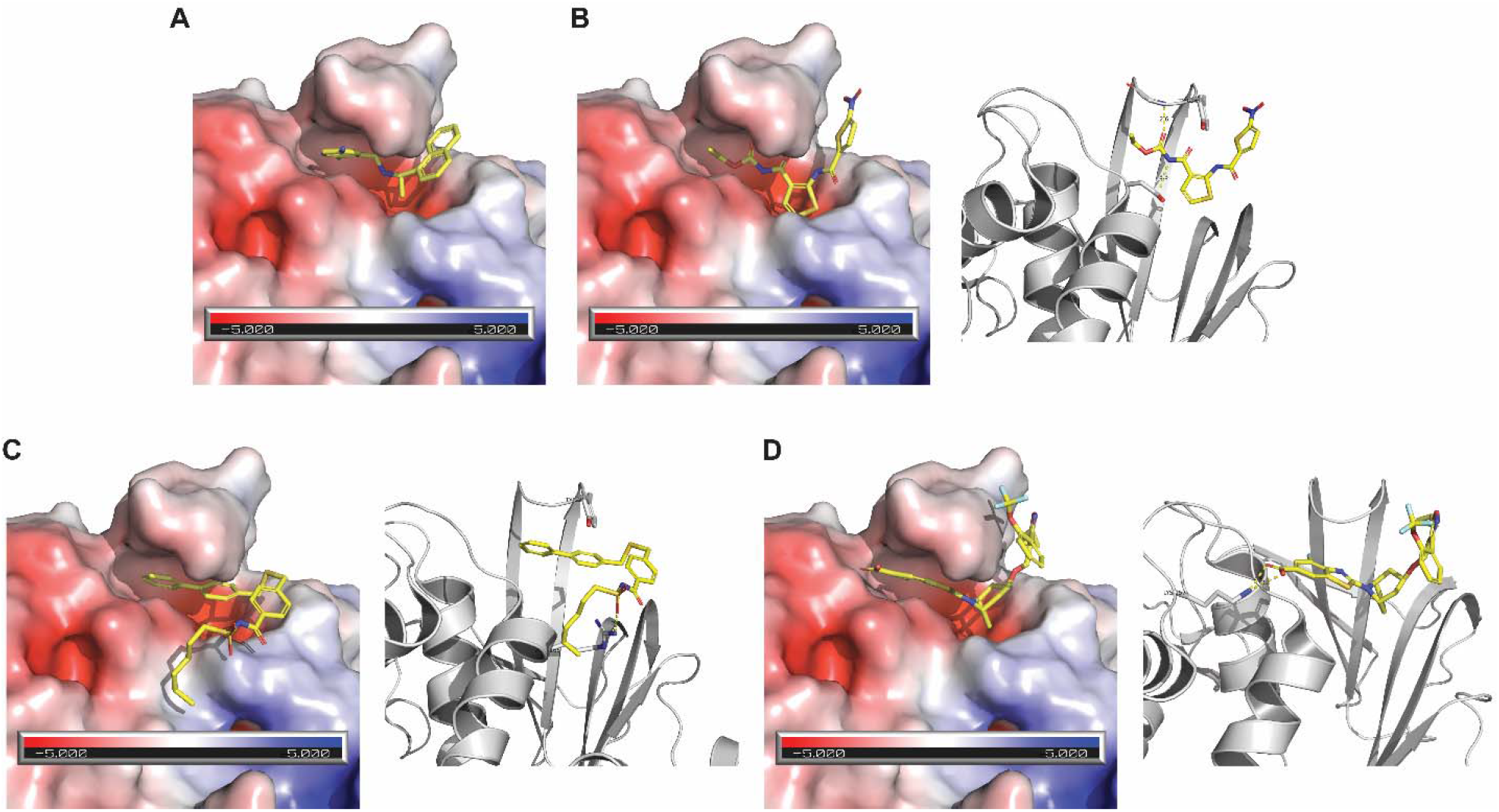
Molecular docking of SARS-CoV-2 PL^pro^ inhibitors GRL0617 (A), EACC(B), KY-226 (C), and tropifexor (D) in PL^pro^ (PDB: 7JRN).

EACC forms two hydrogen bonds with PL^pro^, one from the EACC carbonyl with the Gln269 main chain amide NH, and another from the EACC imide NH with the Asp164 side chain carboxylate (Figure 5B). In addition, the nitro-thienyl ring forms a π-π interaction with the Tyr268 side chain phenol. For KY-226, the benzene ring in the biphenyl substitution similarly forms a π-π interaction with the Tyr268 side chain phenol. The sulfone from KY-226 forms a hydrogen bond with the Arg166 side chain guanidine NH (Figure 5C). The carboxylate from tropifexor forms an ionic bond with the Lys157 side chain ammonium (Figure 5D). The docking poses might provide a guidance for the following lead optimization.

## CONCLUSION

Although PL^pro^ is a validated antiviral drug target, the development of PL^pro^ inhibitors falls behind M^pro^ and RdRp inhibitors. As of date, no PL^pro^ inhibitors have been advanced to the *in vivo* animal model studies yet. The naphthalene compounds such as GRL0617 and its analogs are the only class of validated PL^pro^ inhibitors with antiviral activity against SARS-CoV-2. However, the low metabolic stability of this series of compounds might prevent its further development.^14, 28^ In this study, we aimed to identify structurally novel PL^pro^ inhibitors that can serve as starting points for further optimization. Through screening the MedChemExpress bioactive compound library, three hits EACC, KY-226, and tropifexor were identified as SARS-CoV-2 PL^pro^ inhibitors with IC_50_ values in the single digit micromolar range. Among the three hits, tropifexor appears to be the most promising hit as it also showed potent inhibition against SARS-CoV PL^pro^ (IC_50_ = 5.54 µM) and MERS-CoV PL^pro^ (IC_50_ = 2.32 µM). Tropifexor is a highly potent agonist of the farnesoid X receptor and is currently undergoing phase II clinical trial for nonalcoholic steatohepatitis (NASH) and liver fibrosis.^29^ In addition to the inhibition of PL^pro^ mediated cleavage of viral polyprotein substrate, tropifexor also inhibited the deubiquitination and deISGlation activities of SARS-CoV-2 PL^pro^.

Consistent with the enzymatic inhibition, tropifexor showed dose-dependent stabilization of SARS-CoV-2 PL^pro^ in the thermal shift binding assay. Importantly, tropifexor displayed cellular PL^pro^ inhibitory activity in the FlipGFP assay and the antiviral activity against SARS-CoV-2 in Calu-3 cells. Although the low selectivity index (SI = 6.97) of tropifexor in the antiviral assay prevents its direct repurposing as a SARS-CoV-2 antiviral, the discovery of tropifexor as a novel PL^pro^ inhibitor provides an additional scaffold for further medicinal chemistry optimization.

## MATERIALS AND METHODS

### Protein Expression and Purification

Detailed expression and purification procedures untagged SARS-CoV-2 PL^pro^ and SARS-CoV-2 M^pro^ were described in our previous publications.^15, 30^ SARS-CoV papain-like protease gene (ORF 1ab 1541-1855) (accession # <u>AEA10621.1</u>) from strain SARS coronavirus MA15 with *E. coli* codon optimization in the pET28b-(+) vector was ordered from GenScript. Then the SARS-CoV PL^pro^ gene (ORF 1ab 1541-1855) was subcloned from the pET28b-(+) to pE-SUMO vector according to the manufacturer’s protocol (LifeSensors Inc., Malvern, PA). The forward primer with the Bsa I site is GCGGTCTCAAGGTGAGGTGAAGACCATCAAAGTGTTCACCACC; the reverse primer with a Bsa I site is GCGGTCTCTCTAGATTATTTAATGGTGGTGGTATAGCTGGTTTCCTTGTAG. The expression and purification protocol of SARS-CoV PL^pro^ is identical to SARS CoV-2 PL^pro^.^15^

MERS-CoV PL^pro^ gene (ORF 1ab 1482-1803) (accession # <u>KY581684</u>) from strain MERS coronavirus Hu/UAE_002_2013 with *E. coli* codon optimization in the pET28b-(+) vector was ordered from GenScript. Then MERS-CoV PL^pro^ gene (ORF 1ab 1482-1803) was subclone into pE-SUMO vector with the pair primers: GCGGTCTCAAGGTCAGCTGACCATCGAGGTGCTGGTTACCGTGG and GCGGTCTCTCTAGATTAGTTGCAATCGCTGCTATATTTTTGACCCGGGAAC. The expression and purification protocol of MERS-CoV papain-like protease is identical to SARS CoV-2 PL^pro^.^15^

### FRET substrate synthesis

The SARS-CoV-2 PL^pro^ FRET substrate 1 is Dabcyl-FTLRGG/APTKV(Edans); this substrate was also used as SARS-CoV PL^pro^ and MERS-CoV PL^pro^ substrates. SARS-CoV-2 M^pro^ FRET substrate 2 is Dabcyl-KTSAVLQ/SGFRKME-(Edans). These FRET substrates were synthesized by solid-phase synthesis through iterative cycles of coupling and deprotection using the previously optimized procedure.^31^ Ub-AMC and ISG15-AMC were purchased from BostonBiochem (catalog no. U-550-050 and UL-553-050, respectively).

### Enzymatic Assays

The high-throughput screening was carried out in 384-well format as described previously.^15^ The bioactive compound library consisting of 9,791 compounds was purchased from MedChemExpress (catalog no. HY-L001). The enzymatic reactions for SARS-CoV-2, SARS-CoV, MERS-CoV PL^pro^s were carried out in reaction buffer consisting of 50 mM HEPES pH 7.5, 5 mM DTT and 0.01% Triton X-100. For the IC_50_ measurement with FRET peptide-Edans substrate, the reaction was carried out in 96-well format with 100 µl reaction volume. SARS-CoV-2 PL^pro^ (200 nM) SARS-CoV PL^pro^ (200 nM) or MERS-CoV PL^pro^ (2 µM) was pre-incubated with various concentrations of testing compounds at 30 °C for 30 min before the addition of FRET-peptide substrate to initiate the reaction. The reaction was monitored in a Cytation 5 image reader with filters for excitation at 360/40 nm and emission at 460/40 nm at 30 °C for 1 h. The initial enzymatic reaction velocity was calculated from the initial 10 min enzymatic reaction via a linear regression function and was plotted against the substrate concentrations in Prism 8 with a four-parameter dose-response function. For the IC_50_ measurements with Ub-AMC or ISG15-AMC substrate, the reaction was carried out in 384-well format in 50 µl reaction volume. In the Ub-AMC cleavage assay, the final SARS-CoV-2 PL^pro^ concentration is 50 nM, and substrate Ub-AMC concentration is 2.5 μM. IN the ISG15-AMC assay, the final SARS-CoV-2 PL^pro^ concentration is 2 nM, and substrate ISG15-AMC concentration is 0.5 μM. The SARS-CoV-2 M^pro^ enzymatic assays were carried out in the reaction buffer containing 20 mM HEPES pH 6.5, 120 mM NaCl, 0.4 mM EDTA, 20% glycerol, and 4 mM DTT as described previously.^30, 32^ **Differential Scanning Fluorimetry (DSF)**. The thermal shift binding assay (TSA) was carried out using a Thermo Fisher QuantStudio 5 Real-Time PCR system as described previously.^15, 30^ Briefly, 4 μM SARS-CoV-2 PL^pro^ protein in PL^pro^ reaction buffer (50 mM HEPES pH 7.5, 5 mM DTT and 0.01% Triton X-100) was incubated with various concentrations of testing compounds at 30 °C for 30 min. 1× SYPRO orange dye was added, and the fluorescence of each well was monitored under a temperature gradient range from 20 to 90 °C with 0.05 °C/s incremental step. The melting temperature (Tm) was calculated as the mid-log of the transition phase from the native to the denatured protein using a Boltzmann model in Protein Thermal Shift Software v1.3.

### Cell-Based FlipGFP PL^pro^ Assay

Plasmid pcDNA3-PLpro-flipGFP-T2A-mCherry was constructed from pcDNA3-TEV-flipGFP-T2A-mCherry.^15^ SARS-CoV-2 PL^pro^ expression plasmid pcDNA3.1-SARS2 PL^pro^ was ordered from Genscript (Piscataway NJ) with codon optimization. For transfection, 293T cells were seeded into 96-well Greiner plate (catalog no. 655090) to overnight with 70-90% confluency. 50 ng of pcDNA3-PLPro-flipGFP-T2A-mCherry plasmid and 50 ng of protease expression plasmid pcDNA3.1-PL^pro^ were added to each well in the presence of transfection reagent TransIT-293 (Mirus) according to manufacturer’s protocol. Three hours after transfection, 1 μL of testing compound was added to each well at 100-fold dilution. Images were acquired 2 days after transfection with a Cytation 5 imaging reader (Biotek) GFP and mCherry channels and were analyzed with Gen5 3.10 software (Biotek). SARS-CoV-2 PL^pro^ protease activity was calculated by the ratio of GFP signal over the mCherry signal. The FlipGFP PL^pro^ assay IC_50_ value was determined by plotting the GFP/ mCherry signal over the compound concentration with a four-parameter dose-response function in Prism 8. The mCherry signal alone was utilized to evaluate the transfection efficiency and compound cytotoxicity.

### Molecular modeling of the binding of EACC, KY-226, and tropifexor to SARS-CoV-2 PL^pro^

Docking was performed using Schrödinger Glide standard precision. The SARS-CoV-2 PL^pro^ structure was downloaded from PDB code 7JRN. The final docking poses were generated in PyMOL. The protein electrostatistics surface was generated using the APBS Electrostatistics model in PyMOL.

## AUTHOR INFORMATION

### Author Contributions

J.W. and C.M. conceived and designed the study; C.M. performed the high-throughput screening, enzymatic assays, and thermal shift binding assay. Y. W. and J. C. helped with the protein expression and purification, and the enzymatic assays. J.W. wrote the manuscript with the input from C.M.

## ACKNOWLEDGMENTS

This research was partially supported by the National Institute of Allergy and Infectious Diseasess of Health (NIH) (grants AI147325, AI157046, and AI158775) and the Arizona Biomedical Research Commission Centre Young Investigator grant (ADHS18-198859) to J. W. The SARS-CoV-2 antiviral assay in Calu-3 cells was conducted by Drs. David Schultz and Sara Cherry at the University of Pennsylvania (USA) through the NIAID preclinical service under a non-clinical evaluation agreement.

## Table of Contents Use Only

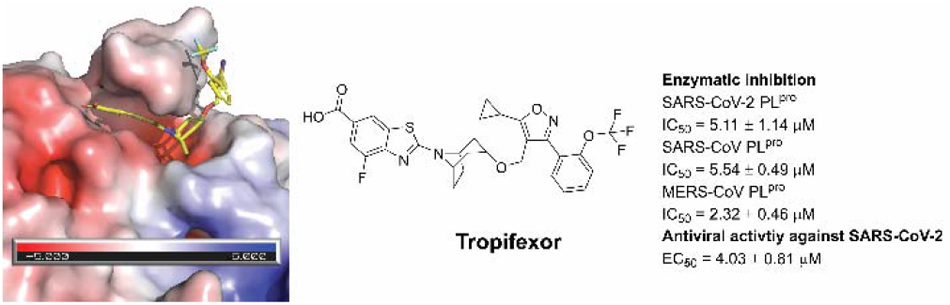

Drug repurposing screening identified tropifexor as a potent SARS-CoV-2 papain-like protease inhibitor with antiviral activity.

## Notes

### Competing Interest Statement

The authors have declared no competing interest.

